# Identification and characterization of a skin microbiome on *Caenorhabditis elegans* suggests environmental microbes confer cuticle protection

**DOI:** 10.1101/2024.02.21.581412

**Authors:** Nadia Haghani, Robert H. Lampe, Buck Samuel, Sreekanth H. Chalasani, Molly A. Matty

**Author notes:** Address Correspondence to: Molly A. Matty. Nadia Haghani, Department of Biology, Stanford University, Palo Alto CA. Molly A. Matty, Department of Biology, University of Portland, Portland OR.

## Abstract

In the wild, *C. elegans* are emersed in environments teeming with a veritable menagerie of microorganisms. The *C. elegans* cuticular surface serves as a barrier and first point of contact with this microbial milieux. Here, we identify microbes from *C. elegans* natural habitats that associate with its cuticle, constituting a simple ‘skin microbiome’. We rear our animals on a modified CeMbio, mCeMbio, a consortium of ecologically relevant microbes. We first combine standard microbiological methods with an adapted micro skin-swabbing tool to describe the skin-resident bacteria on the *C. elegans* surface. Further, we conduct 16S rRNA gene sequencing studies to identify relative shifts in the proportion of mCeMbio bacteria upon surface-sterilization, implying distinct skin-and gut-microbiomes. We find that some strains of bacteria, including *Enterobacter sp.* JUb101, are primarily found on the nematode skin, while others like *Stenotrophomonas indicatrix* JUb19 and *Ochrobactrum vermis* MYb71 are predominantly in the animal’s gut. Finally, we show that this skin microbiome promotes host cuticle integrity in harsh environments. Together, we identify a skin microbiome for the well-studied nematode model and propose its value in conferring host fitness advantages in naturalized contexts.

**Importance:** The genetic model organism *C. elegans* has recently emerged as a tool for understanding host-microbiome interactions. Nearly all of these studies either focus on pathogenic or gut-resident microbes. Little is known about the existence of native, non-pathogenic skin microbes or their function. We show that members of a modified *C. elegans* model microbiome, mCeMbio, can adhere to the animal’s cuticle and confer protection from noxious environments. We combine a novel micro-swab tool, the first 16S microbial sequencing data from relatively unperturbed *C. elegans*, and physiological assays to demonstrate microbially mediated protection of the skin. This work serves as a foundation to explore wild *C. elegans* skin microbiomes and use *C. elegans* as a model for skin research.

## Introduction

All animals are in contact with microbial communities, termed the microbiome. Microbiomes play a large role in shaping host physiology, adaptation, fitness, and even behavior(De Palma et al. 2015; Baurecht et al. 2018; Yildirim et al. 2010; Sampson and Mazmanian 2015). Organisms and their microbial inhabitants share their first interactions at the integument of the host (Naik et al. 2015). This enveloping layer, whether it be skin, membranes, or a cuticle, plays an essential role in host-microbe relationships, primarily as a barrier to promote host survival. These essential roles coupled with dynamic microbial communities provide a valuable interface to study host-microbe interactions. Emerging studies specifically focus on the effects of single, pathogenic bacteria, which are often localized to the gut, on animal behavior and physiology (O’Donnell et al. 2020; Chiu et al. 2013; Y. Zhang, Lu, and Bargmann 2005; Gill et al. 2006). Although single species studies allow a thorough investigation of host-microbe interactions, dominant microbial species do not fully recapitulate entire communities of microbes (Charisse Petersen and Round 2014). Moreover, a majority of these studies focus on pathogen invasion mechanisms of the skin, or pathogen-commensal microbial interactions (Ramsey et al. 2016), whereas host-commensal dynamics at the skin surface remain impactful but understudied (Naik et al. 2015; 2012).

This is also the case for the soil and fruit-dwelling nematode, *Caenorhabditis elegans*. Despite the subtle implication of surface-adherent bacteria, host-microbe studies at the surface layer are often focused on pathogens (Jansson 1994; J. Hodgkin, Kuwabara, and Corneliussen 2000; Darby et al. 2007). In the wild, *C. elegans* are found in rotting organic matter filled with microorganisms (Samuel et al. 2016; Dirksen et al. 2016; Berg et al. 2016). Therefore, interaction between external microbes and the worm’s integument—the cuticle—is inevitable and likely a dynamic process (Félix and Duveau 2012), Moreover, these nematodes rely on their cuticle health to protect against environmental stressors such as harmful toxins (Kucharíková et al. 2023), desiccation (Erkut et al. 2013), or pathogens (Sellegounder et al. 2019; Forman-Rubinsky, Cohen, and Sundaram 2017). The function of the *C. elegans* cuticle and its many structures have been explored using mutants in the underlying genes. Animals with defective cuticle structures display a number of phenotypes including motor abnormalities, a higher susceptibility to chemically induced cuticle degradation, and protection from skin pathogens (Levy, Yang, and Kramer 1993; Partridge et al. 2008; Sandhu et al. 2021; Höflich et al. 2004). Several modes of fungal and single-bacterial adherence and pathogenesis have been characterized (Jansson 1994; J. Hodgkin, Kuwabara, and Corneliussen 2000; Darby et al. 2007) including *Leucobacter* strains known to externally invade *C. elegans* surfaces (J. Hodgkin, Kuwabara, and Corneliussen 2000; Muir and Tan 2008). However, interactions involving ecologically relevant commensals, and moreover, communities of microbes that represent wild *C. elegans* encounters, remain underexplored.

Nematodes are maintained on bacterial lawns in laboratory settings. Together with their genetic tools and transparent bodies, *C. elegans* have recently gained appeal in microbiome studies (F. Zhang et al. 2017; Frézal and Félix, n.d.). In the laboratory setting, a single bacterium *Escherichia coli* (usually a single strain, OP50) serves as the animal’s food source and external environment and minimally colonizes the *C. elegans* intestine for most of its life (Podshivalova, Kerr, and Kenyon 2017). However, *E. coli* is not a substitute for a microbiome of *C. elegans*, nor is it extensively associated with the nematode in the wild. A recent meta-analysis of the *C. elegans* intestinal microbiota produced a comprehensive analysis of the host’s natural gut inhabitants (Samuel et al. 2016; Dirksen et al. 2016; Berg et al. 2016). This collaborative effort prompted the creation of the *C. elegans* Microbiome (CeMbio), a model microbiome consisting of the 12 most abundant gut bacteria families naturally associated with *C. elegans* (Dirksen et al. 2020, 202). Individual isolates from this consortium are known to affect growth, food choice behavior, and physiology, as well neural degradation through metabolite secretion (Chen et al. 2022; Kohar Annie B. Kissoyan et al. 2022; Urquiza-Zurich et al. 2023; Carola Petersen et al. 2021; Chai et al. 2024). However, these studies only consider the influence and composition of CeMbio bacteria in the gut and have not examined their potential impact at other body sites like the cuticle.

In preparation for intestinal microbial sequencing, animals are traditionally subjected to harsh bleach washes and surface-sterilization protocols to remove cuticle-resident bacteria (Palominos and Calixto 2020). Moreover, the first study using CeMbio intensified previously established washing steps to more effectively remove bacteria that were hypothesized to still remain on the surface, even following multiple washing steps (Dirksen et al. 2020). The examination of nematode microbiomes in the absence of stringent surface bleach sterilization may provide worthwhile insights into the effects of a comprehensive, complex, and natural microbiome on cuticle physiology and host biology.

To investigate the persistence, existence, and composition of a cuticle-resident microbiome, we develop a novel micro-swabbing tool in combination with bacterial growth curves to find that animals reared on a complex microbiome harbor cuticle-resident bacteria. To supplement this finding, we enumerate the abundance of bacteria on surface-sterilized animals to demonstrate that surface-sterilization protocols remove a significant number of skin-resident or associated bacteria. We then conduct 16S rRNA gene sequencing to characterize a skin microbiome on animals reared on complex microbiomes similar to CeMbio, which we call modified CeMbio (mCeMbio, Table 1). Importantly, we find that the microbiome of non-sterilized *C. elegans* is microbially distinct from the microbiome of the *C. elegans* gut and the surrounding environment. We identify skin-and gut-dominant isolates of this microbial consortium that demonstrate distinct attachment strengths to the cuticle. Finally, we show that colonization with mCeMbio and with several mCeMbio isolates confers protection for the host against noxious stimuli.

**Table 1:** Bacterial strains used in this study.

| ID | Genus | Species | Source |
| --- | --- | --- | --- |
| JUb19 | *Stenotrophomonas* | *maltophilia* | mCeMbio, CGC CeMbio |
| JUb44 | *Chryseobacterium* | *scophthalmum* | mCeMbio, CGC CeMbio |
| JUb101 | *Enterobacter* | *sp.* | mCeMbio |
| JUb134 | *Sphingomonas* | *molluscorum* | mCeMbio, CGC CeMbio |
| MYb10 | *Acinetobacter* | *guillouiae* | mCeMbio, CGC CeMbio |
| MYb11 | *Pseudomonas* | *lurida* | mCeMbio, CGC CeMbio |
| MYb71 | *Ochrobactrum* | *pecoris* | mCeMbio, CGC CeMbio |
| BIGb0170 | *Sphingobacterium* | *multivorum* | mCeMbio, CGC CeMbio |
| BIGb0172 | *Comamonas* | *piscis* | mCeMbio, CGC CeMbio |
| BIGb0393 | *Pantoea* | *nemavictus* | mCeMbio, CGC CeMbio |
| MSPm1 | *Pseudomonas* | *mendocina* | mCeMbio, CGC CeMbio |
| CEN2ent1 | *Enterobacter* | *xiangfangensis* | mCeMbio, CGC CeMbio |
| CBX151 | *Leucobacter* | *celer subsp. astrifaciens* | Known skin bacterium |
| OP50 | *Escherichia* | *coli* | Standard food |

## Results

### A skin swab protocol reveals that abundant mCeMbio bacteria persist on C. elegans **surface**

We sought to directly identify the existence of a resident skin microbiome on the *C. elegans* cuticle. However, existing microbiological techniques in the nematode are designed to isolate the entirety of the animal’s microbiota rather than skin microbes. We postulated that the entire microbiota consists of gut-resident and surface-resident microbes, and that these parts can be separated with varying intensities of surface-washing steps (Figure 1A). For instance, surface-sterilization, hereafter referred to as “bleaching”, removes all skin bacteria and isolates gut-resident bacteria (Figure 1A). Moreover, we hypothesize that animals with more skin-resident bacteria will demonstrate larger decreases in bacterial counts following bleaching. Using a Colony Forming Unit (CFU) assay, we assessed the existence of surface-adherent microbes on animals reared on OP50 or mCeMbio. First, we observe greater colonization of animals reared on mCeMbio compared to OP50 (Figure 1B). We also observe no significant difference between the number of CFUs from whole animal lysates that were serially washed or bleached after being reared on OP50 (<10^3^ bacteria per worm, Figure 1B). However, we observe a significant decrease in the number of CFUs from whole animals after bleaching (<10^4^ bacteria per worm) compared to serially washed animals (>10^6^ bacteria per worm). From these data, we find that animals reared on mCeMbio, but not OP50, harbor skin-resident bacteria. Moreover, to account for free-living bacteria that are not in contact with the skin, we analyzed the CFUs from supernatants after serial washes. We observe that there are OP50 and mCeMbio bacteria present in the supernatant after three washes, which might reflect transient gut or skin microbes that have been shed and highlight the limited extent of these methods for identifying skin-resident microbes.

**Figure 1:**
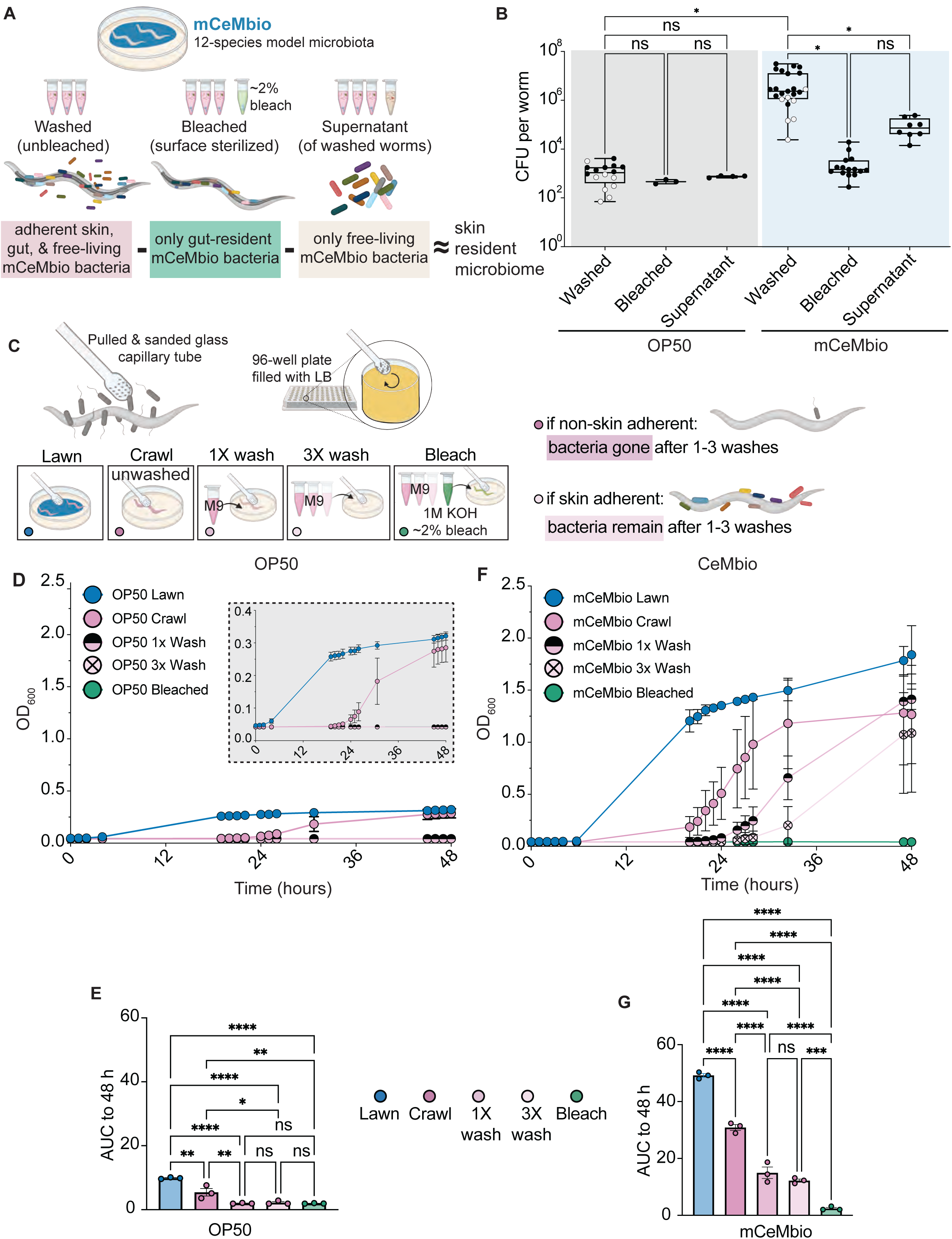
CFU analysis and novel skin swab protocols reveal that abundant bacteria persist on *C. elegans* surface. A) Diagram showing analysis of CFUs from washed, bleached, and liquid supernatants roughly estimates skin-resident bacterial abundance. B) Average CFUs per worm in each sample type. Each dot represents an experiment with 50 worms. White dots represent experiments conducted without a bleached counterpart. Lower quartile, mean, and upper quartile marked by box and whisker plot. C) Diagram showing skin swabbing and washing procedures used throughout Figure 1 and Figure 3. D, F) OD_600_ of swabbed bacteria from animals reared on OP50 (D) and modified CeMbio (mCeMbio) (F) is measured for 72 hours. Data shown are representative of average OD600 values from an individual experiment with n≥5 animals for each condition. Mean with standard deviation in bars. E, G) Mean area under the curve (AUC) for swabs of animals raised on OP50 (E) or mCeMbio (G) from three independent experiments, with each dot representing the average AUC for an experiment measured to 48 hours post swab. Additional experiments are found in Supplementary Figure 1 A-D. B) Brown-Forsythe One-Way ANOVA tests (for unequal SDs) with Dunnett’s T3 for multiple comparisons test. E, G: Ordinary 1-way ANOVA with Tukey’s multiple comparisons test. Error bars are SEM. * P < 0.05, ** P <0.01, *** P<0.001, **** P <0.0001

Inspired by existing methods to identify skin microbes (Grice et al. 2009; Jani and Briggs 2014), we created a microscopic nematode-sized skin swab. This reusable glass tool is swabbed across the length of each animal (Figure 1C). To eliminate non-adherent, free-living microbes and enrich for putative skin-resident microbes, we compare unwashed animals (‘crawl’) to animals that have been washed once or three times. As a negative control, we include animals that have been bleached. Swabs from animals reared on OP50 indicate that OP50 grows from lawns and unwashed animals, but even a single wash eliminates all skin-associated OP50 bacteria (Figure 1D, Supplementary Figure 1 A, C). Specifically, growth curves of OP50 from singly (1X) and triply (3X) washed animals are indistinguishable from the swabs from bleached animals (Figure 1E). Swabs from animals reared on mCeMbio indicate that viable bacteria can be isolated from lawns and unwashed animals as well as animals that have been washed up to three times (Figure 1F, Supplementary Figure 1B, D). Growth curves of mCeMbio from animals washed up to three times had ∼7-fold higher levels of bacteria compared to bleached animals (Figure 1G). These findings suggest that there are significant populations of adherent mCeMbio bacteria on the *C. elegans* cuticle that are removed by standard bleach-based surface-sterilization protocols.

### Surface sterilization alters the relative proportion of mCeMbio bacteria, suggesting distinct surface-resident communities

We sought to define the species of mCeMbio bacteria that reside on the *C. elegans* cuticle and determine whether they are distinct from the microbiota of the gut and the surrounding environment. We performed 16S rRNA gene sequencing on bleached *C. elegans*, serially washed *C. elegans*, and their bacterial lawns. In two separate experiments, we find that the bacterial communities of bleached animals have a microbiota that is distinct from animals that have been serially washed. In bleached animals, we observed significantly higher levels of *Stenotrophomonas indicatrix* JUb19 and *Ochrobactrum vermis* MYb71, while unbleached animals contain more *Enterobacter sp.* JUb101 and *Sphingobacterium multivorum* BIGb0170 (Figure 2A-C). Despite significant differences between the relative abundance of bacteria across experiments 1 and 2 (Supplementary Figure 2A-C), we observe consistent differences in microbiota composition of bleached and serially washed animals, both of which are distinct from bacterial lawns (Figure 2E, F). These data suggest that bacterial communities of bleached animals are significantly different than those of serially washed animals by Bray-Curtis distances (PERMANOVA, Figure 2G, I). Furthermore, within sample alpha diversity, expressed as the Shannon index, of each treatment group is similar, but mCeMbio bacterial lawns display greater diversity than bleached or serially washed *C. elegans* in experiment 2 (Figure 2H, J). Collectively, we suggest that the microbiota of *C. elegans* with an intact skin microbiota is distinct from the microbiota of the *C. elegans* gut, and further distinct from the environment on which the animals were raised. Specifically, we hypothesize that more abundant species in bleached animals (JUb19 and MYb71) are more gut-specific whereas species more abundant in serially washed animals (JUb101 and BIGb0170) are more skin-specific.

**Figure 2:**
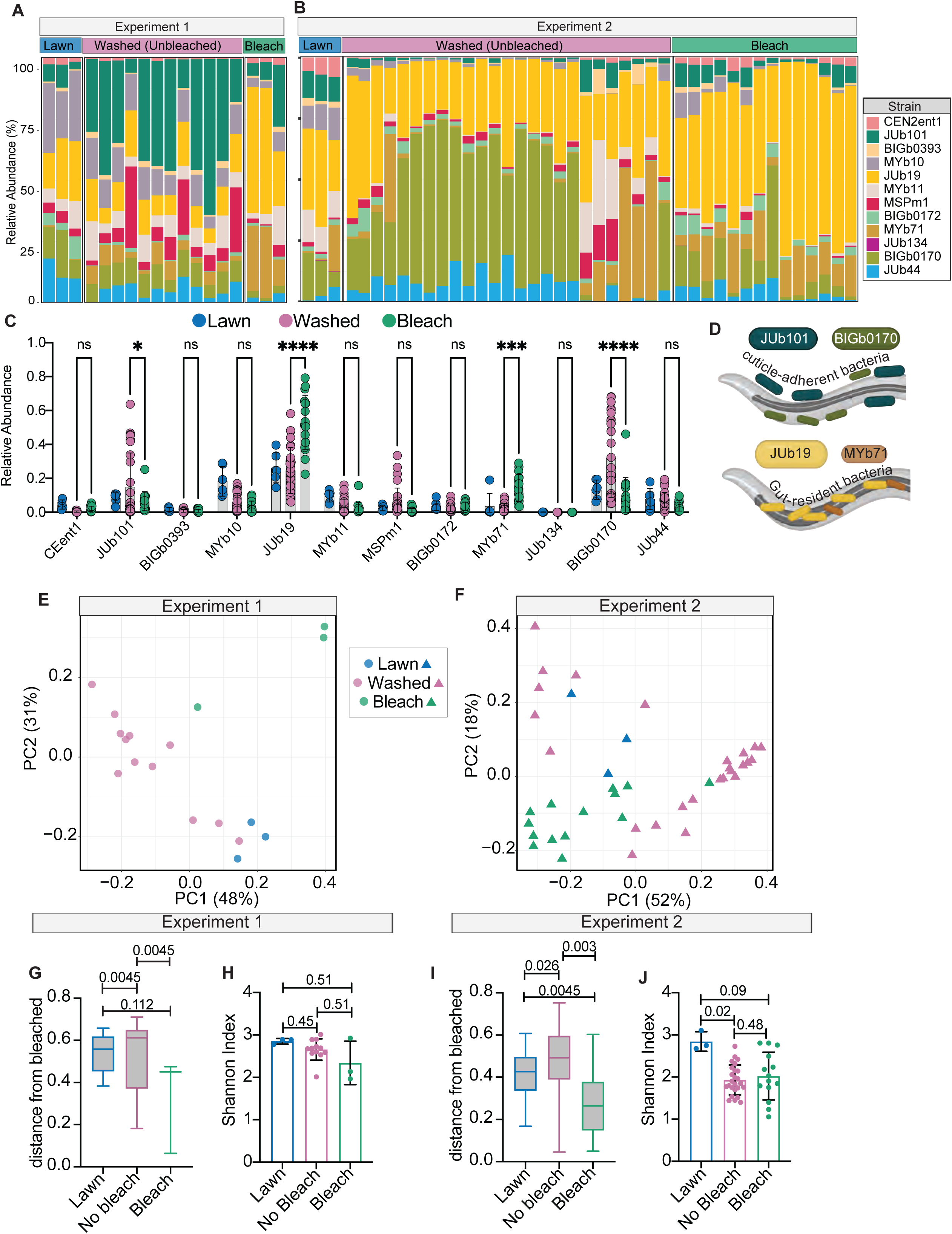
Surface sterilization alters the relative proportion of mCeMbio bacteria. A-B) Proportion of reads from mCeMbio bacteria in bacterial lawns, washed (unbleached) and bleached animals from experiments 1 (A) and 2 (B). C) Mean proportion of reads for lawns (blue), washed (pink) and bleached (green) N2 animals, combined from matched bleach/washed sets in experiments 1 and 2. Each dot represents one plate of animals. D) Diagram of bacterial strains of interest and their expected niches, not to scale. E, F) Principal coordinate analysis of Bray-Curtis dissimilarities for samples from experiment 1 (E) and experiment 2 (F). C) 2-way ANOVA – significant difference between bacteria and the interaction between bacteria and bleach/no bleach groups (P <0.0001) with “Šídák’s multiple comparisons test” within each bacteria strain. G, I) Bray-Curtis dissimilarities as distance from the bleached group for experiment 1 (G) and experiment 2 (I) with Pairwise PERMANOVA q-values. H, J) Shannon index for each treatment in experiment 1 (H) and 2 (J) with pairwise Kruskal-Wallis q-values. * P < 0.05 by, ** P <0.01, *** P<0.001, **** P <0.0001

### Specific mCeMbio microbes associate with the C. elegans cuticle

Informed by 16S relative abundance data, we sought to identify the effects of bleaching or serial washes on animals reared on JUb19 or JUb101. JUb101 is the only predominantly skin-resident microbe on which *C. elegans* develop at similar rates to OP50 and mCeMbio (Dirksen et al. 2020 and our observations) and JUb19 is the gut-resident microbe with the highest relative abundance (Figure 2C).

As in Figure 1, we performed CFU analysis on our microbes of interest. We observe more CFUs from lysates of serially washed animals reared on JUb101 (>10^6^ bacteria per worm) than lysates of serially washed animals reared on JUb19 (<10^5^ bacteria per worm) (Figure 3A). There is no statistically significant difference between the number of CFUs from lysates of serially washed or bleached animals reared on JUb19, although JUb19 is still found on the surface. This is indicated by ∼10^4^ CFU per worm found in and on serially washed animals, compared to the ∼10^2^ CFU per worm in bleached animals (Figure 3A). Conversely, in animals raised on JUb101, there is a statistically significant decrease in the number of CFUs from whole animals after bleaching (∼10^2^ bacteria per worm) compared to serially washed animals (>10^6^ bacteria per worm), indicating the presence of JUb101 as a persistent and abundant skin-resident microbe that is removed with surface sterilization protocols but not serial washes (Figure 3A). To account for free-living bacteria that are loosely attached to the cuticle or transient within the worm, we analyzed the CFUs from supernatants following serial washes. We observe that JUb101 and JUb19 are both present in these supernatants, but in lower abundance than in washed groups.

**Figure 3:**
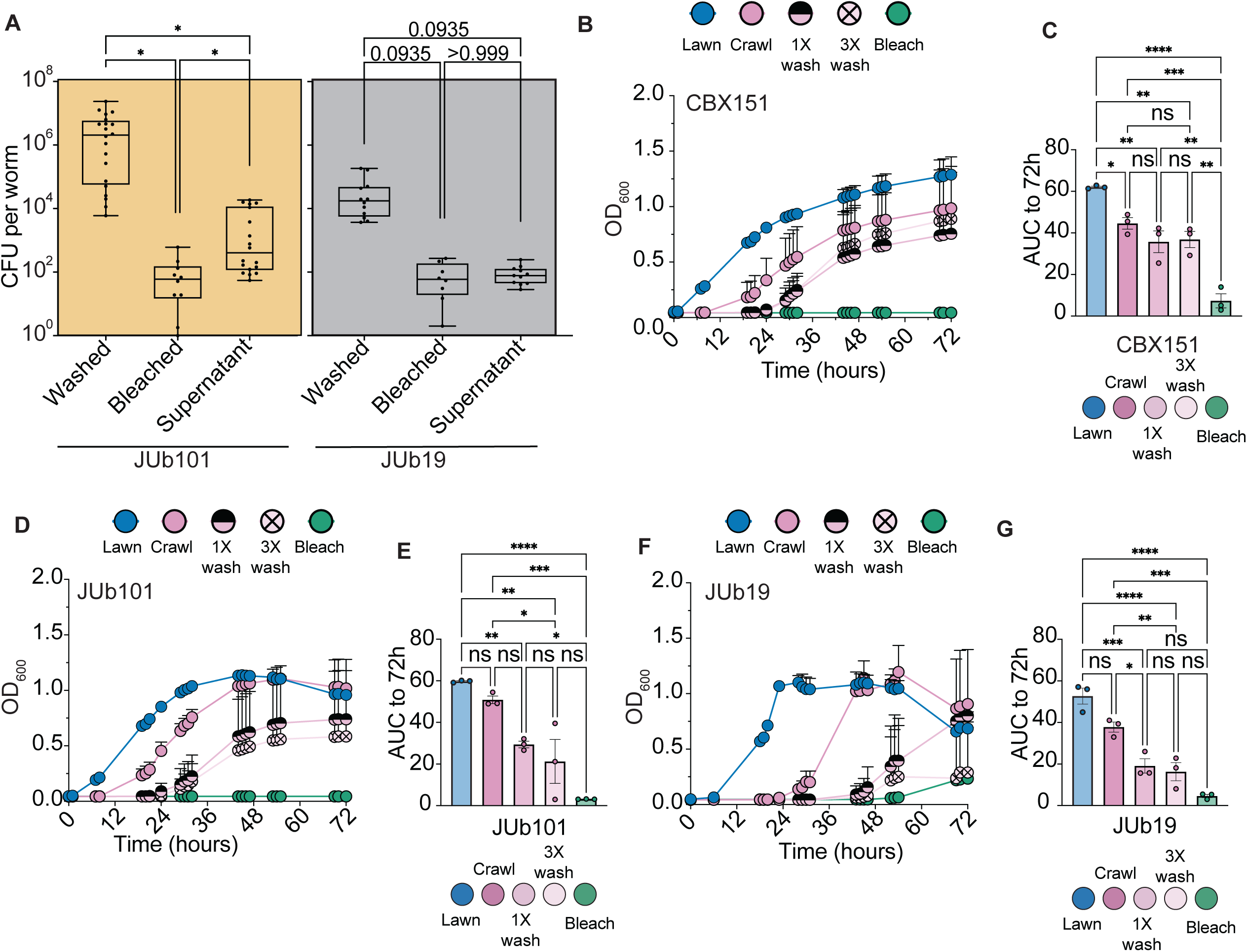
JUb19 and JUb101 display variable strengths of cuticle attachment. A) Average Colony Forming Units per worm in each sample type. Each dot represents an experiment with 50 worms. Lower quartile, mean, and upper quartile marked by box and whisker plot. B,D,F) OD_600_ of swabbed bacteria from animals reared on CBX151 (B), JUb101 (D), and JUb19 (F) is measured for 72 hours. Data shown is representative of average OD_600_ values from an individual experiment with n≥5 animals for each condition. Mean with standard deviation bars displayed. C,E,G) Mean area under the curve (AUC) for swabs of animals raised on CBX151 (C), JUb101 (E) or JUb19 (G) from three independent experiments, with each dot representing the average AUC for an experiment to 72 hours post swab. Additional experiments are found in Supplementary Figure 3 A-F. A) Brown-Forsythe One-Way ANOVA with Dunnett’s T3 Multiple Comparison’s Test C, E, G: Ordinary 1-way ANOVA with Tukey’s multiple comparisons test. Error bars are SEM. * P<0.05, ** P<0.05, ***P<0.001 **** P <0.0001.

To directly compare JUb19 and JUb101 as putative gut and skin-resident microbes, respectively, we swabbed animals reared on JUb19, JUb101, or CBX151. We include CBX151 (*Leucobacter celer* subsp*. astrifaciens*, Verde1) as a control bacterium that is known to strongly adhere to the *C. elegans* cuticle (Clark and Hodgkin 2015). We hypothesized that growth curves of swabbed skin-resident bacteria should more closely resemble those of CBX151 and mCeMbio animals (Figure 1G), whereas growth curves of swabbed gut-resident bacteria should resemble OP50 (Figure 1E). As expected, swabs from animals reared on CBX151 grow regardless of the number of washes (Figure 3B). After 72 hours of growth, there is no discernable difference between swabs from animals that were unwashed (‘crawl’), singly, or triply washed (Figure 3C, Supplementary Figure 3A, B). Similarly, swabs from animals reared on JUb101 grow regardless of the number of washes (Figure 3D). After 72 hours of growth, swabs from JUb101-reared animals that have been singly washed are not distinct from unwashed animals, but are distinct from bleached animals (Figure 3E, Supplementary Figure 3C, D). Conversely, swabs from animals reared on JUb19 indicate that bacterial growth from unwashed ‘crawl’ animals is distinct from singly and triply-washed animals (Figure 3F, G). Swabs from singly and triply-washed JUb19 animals are indistinguishable from swabs of bleached animals (Figure 3G, Supplementary Figure 3E, F). Together, JUb101 growth curves resemble those of CBX151, suggesting that JUb101 is also likely a cuticle-resident microbe.

### CeMbio bacteria promote wildtype and mutant C. elegans cuticle integrity

Given that mCeMbio contains both skin-and gut-resident bacteria, with variable strengths of attachment on the nematode cuticle, we sought to observe a potential role for skin-adherent bacteria. We hypothesized that bacteria associated with the cuticle (JUb101, BIGb0170, and CBX151 as a known skin-adherent control) would protect animals from noxious environments more than gut-associated ones (JUb19, MYb71). To test this, we performed a cuticle resistance assay using treatment of animals with a harsh (5%) bleach solution (Figure 4A). In this assay, the time to burst is a readout of cuticle integrity and a potential indication of the protective roles that bacteria may have on worm physiology (Gravato-Nobre et al. 2005; Loer et al. 2015). We found that wild-type animals reared on CBX151 and mCeMbio community take significantly longer to burst compared to OP50 controls and therefore exhibit greater cuticle resistance (Figure 4B). Animals were also reared on mCeMbio isolates individually, and we observed a range impact on cuticle integrity. Some bacterial species, including gut-resident *Ochrobactrum* MYb71, significantly promoted cuticle integrity with longer times to burst. Other bacteria, including the skin-associated JUb101 and gut-associated JUb19, did not alter animal burst times compared to OP50. *Sphingomonas* JUb134 was the only mCeMbio strain to significantly decrease host cuticle integrity, further supporting that most mCeMbio strains are non-pathogenic in this context (Dirksen et al. 2020). However, neither JUb101 nor JUb19 consistently change *C. elegans* cuticle burst time on their own (Figure 4B). This suggests that interspecies interactions present in mCeMbio may promote cuticle protection or the N2 genotype lacks sensitivity in this assay and masks the subtle impact on the cuticle.

**Figure 4:**
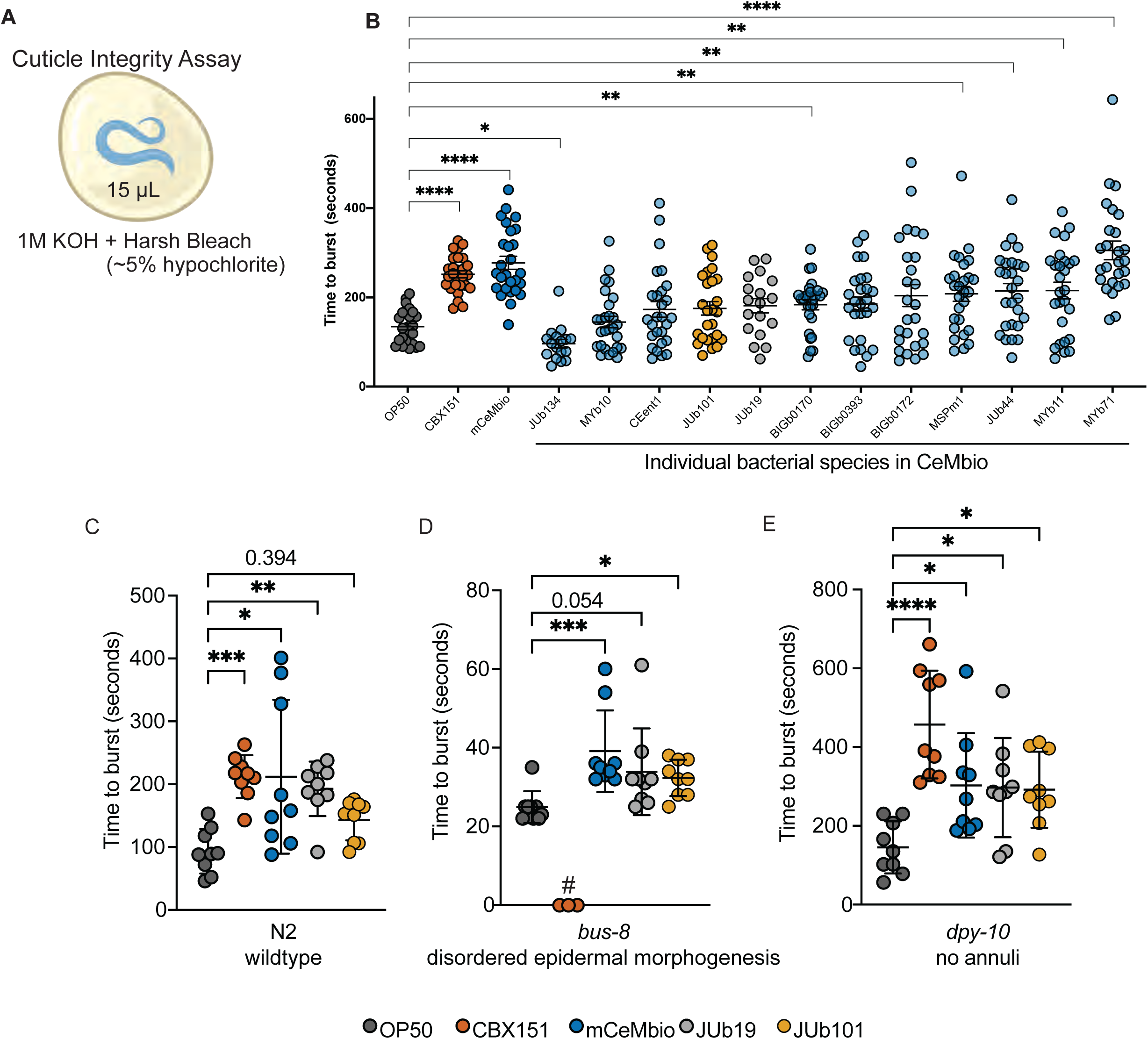
CBX151, mCeMbio and some individual species protect wildtype and mutant *C. elegans* from harsh bleach. A) Schematic of the bleach sensitivity assay using harsh bleach. B) The average time to burst for N2 day 1 adult worms raised on CBX151, mCeMbio, and each individual mCeMbio strain. Each dot represents an individual animal. Mean with SEM. n=9 animals per experiment, N=3 experiments. C-E) Representative data for time to burst for N2 (C), *bus-8* (D), or *dpy-10* (E) animals raised on OP50, CBX151, mCeMbio, JUb19, and JUb101. Each dot represents an individual animal n=9, mean with SD. All experimental averages are in Table 2 and normalized data from multiple experiments are in Supplementary Figure 4 B-D. B) Brown-Forsythe One-Way ANOVA with Dunnett’s T3 Multiple Comparisons Test. C-D) Kruskal-Wallis One-Way ANOVA for unequal SDs with Dunn’s multiple comparison test E) Ordinary One-Way ANOVA with Dunnett’s Multiple Comparison Test. * p<0.05, ** p<0.01, *** p<0.001, **** p<0.0001. #CBX151 was unable to be tested in *bus-8* mutants

**Table 2:** All cuticle burst times. Average time to cuticle burst for 9 animals from each genotype (N2, *bus-8,* and *dpy-10*) raised on OP50, CBX151, mCeMbio, JUb19 or JUb101. Set 1 is shown in Figure 4C-E. All data shown in Supplementary Figures B, C, and D

|  | Bacteria | Set 1 (Figure 4 C, D, E) | Set 2 | Set 3 | Set 4 | Set 5 | Set 6 |
| --- | --- | --- | --- | --- | --- | --- | --- |
| **N2** | OP50 | 93±35 | 81±28 | 67±31 | 121±38 | 265±112 | 112±30 |
|  | CBX151 | 212±34 | 243±67 | 269±45 | 220±28 | na | na |
|  | mCeMbio | 212±122 | 322±194 | 354±100 | 346±132 | na | na |
|  | JUb19 | 193±43 | 129±47 | 124±42 | 81±41 | na | na |
|  | JUb101 | 143±32 | na | na | na | 218±49 | 187±100 |
| ***bus-8*** | OP50 | 25±4 | 29±3 | 28±3 | 32±5 | 30±7 | 30±6 |
|  | CBX151 | unable to test |  |  |  |  |  |
|  | mCeMbio | 39±10 | 37±8 | 38±7 | 50±21 | na | na |
|  | JUb19 | 34±11 | 42±5 | 31±4 | 39±11 | na | na |
|  | JUb101 | 32±5 | na | na | na | 34±11 | 26±4 |
| ***dpy-10*** | OP50 | 145±66 | 151±105 | 178±117 | 90±48 | 242±159 | 193±123 |
|  | CBX151 | 409±157 | 481±141 | 541±86 | 486±143 | na | na |
|  | mCeMbio | 303±132 | 242±51 | 216±141 | 396±117 | na | na |
|  | JUb19 | 297±126 | 143±82 | 289±158 | 296±139 | na | na |
|  | JUb101 | 280±102 | na | na | na | 504±193 | 391±176 |

To test this, we assessed how these bacteria affect animals with disrupted cuticle structures, thereby sensitizing the assay. We tested animals with mutations in cuticle-related genes: *bus-8, dpy-8, srf-3,* and *dpy-10* (Supplementary Figure 4A, Supplementary Table 1). All genes are known to affect cuticle integrity. *Bus-8* is predicted to encode a glycosyltransferase, enabling proper cuticle molts, and leading to disordered epidermal morphogenesis in strong mutants (Partridge et al. 2008; Gravato-Nobre et al. 2005). *Dpy-8* and *dpy-10* are collagen genes that, upon disruption, cause loss of cuticular annuli structures (Sandhu et al. 2021). *Srf-3* encodes a nucleotide sugar transporter, and mutants have altered cuticle surface compositions that confer biofilm resistance (Höflich et al. 2004). Previous studies have demonstrated *bus-8(e2698)* mutants as more sensitive (Gravato-Nobre et al. 2005) in the cuticle resistance assay, while *dpy-10* mutants have been shown to be more stiff than wildtype animals (Fechner et al., n.d.) and express the innate immune gene *nlp-29* at high levels constitutively(Zugasti et al. 2014) while also being more permeable to a Hoescht staining protocol (Sandhu et al. 2021), suggesting a multifaceted role for *dpy-10* in protection and permeability. Cuticle burst times of OP50-reared *bus-8* mutants are significantly shorter (time to burst = 25 seconds) than of N2 animals (time to burst = 93 seconds), as previously observed in literature (Gravato-Nobre et al. 2005) (Figure 4D, C Supplementary Figure 4A). Despite a disordered epidermal morphology and disrupted cuticle integrity in *bus-8* animals, mCeMbio consistently promotes cuticle integrity (time to burst = 39 seconds) compared to OP50 (time to burst = 25 seconds) (Figure 4D). To normalize across multiple experiments and variable times to burst, we evaluated the fold change in burst time compared to animals raised on OP50. We observed significant differences in the fold change in burst time of *bus-8* animals raised on mCeMbio and JUb19 compared to animals raised on OP50, with mCeMbio providing 1.4-fold increases in burst time and JUb19 providing a 1.3-fold increase; JUb101 does not significantly increase the fold change in burst time but shows variability between experiments (Supplementary Figure 4B-D). Interestingly, the cuticle-adherent CBX151 caused either parental death or suppressed growth in *bus-8* animals, preventing us from testing this condition consistently in this cuticle mutant background. Cuticle burst times of OP50-reared *dpy-10* mutants are significantly longer (time to burst = 145 seconds) than of N2 animals (time to burst = 93 seconds) (Figure 4C, E). *Dpy-10* animals raised on CBX151 (time to burst =457 seconds), mCeMbio (time to burst =302 seconds), JUb19 (time to burst =219 seconds), or JUb101 (time to burst =291seconds) display noticeably longer cuticle burst times compared to *dpy-10* animals raised on OP50 (time to burst = 145 seconds) (Figure 4E). This relationship is consistent with the fold change over OP50 observed for animals raised on CBX151 (3.9-fold), mCeMbio (2.3-fold), JUb19 (1.9-fold), and JUb101 (2.2-fold) (Supplementary Figure 4D). Given the consistency of increased cuticle integrity in both wild-type and cuticle mutants reared on mCeMbio compared to OP50, bacteria-bacteria interactions likely impact *C. elegans* bleach sensitivity. Moreover, *dpy-10* mutants reared on JUb101 or JUb19 promote cuticle protection compared to OP50, unmasking a subtle yet consistent impact these bacteria have on the cuticle. Despite being a single species, CBX151 is also a consistent promoter of cuticle integrity, suggesting a role for adherence to cuticle structures still present on *dpy-10* and *bus-8* mutants.

## Discussion

Bacteria-host interactions are complex and known to impact many aspects of host biology. However, the study of ecologically relevant bacterial communities is often traded for the simplicity of single species. In this study, we demonstrate the importance of ecologically-relevant communities of bacteria and uncover a *C. elegans* skin-resident microbiome. We reared animals on mCeMbio, a consortium of ecologically relevant bacteria, and discovered that most of the mCeMbio bacteria can be removed by bleaching protocols, suggesting that these bacteria may reside on the cuticle surface (Fig 1A, B). We observed that our surface-sterilized worm CFU counts match previous findings (Dirksen et al. 2020) whereas serially washed, unbleached worms have significantly more bacteria (Figure 1B, 3A). We conjectured that there are abundant skin-resident bacteria, approximating counts from simplified arithmetic of [# unbleached CFU] – [# bleached CFU] – [# supernatant CFU] (Figure 1A). To supplement this finding, we developed and implemented a novel micro-swab to isolate skin microbes from *C. elegans* cuticles. We measured the growth of swabbed OP50 and mCeMbio bacteria from *C. elegans* cuticles and observed distinct and repeatable patterns of relative growth between washing conditions (Figure 1D-G); more washes are required to remove mCeMbio bacteria from the cuticle, implying that mCeMbio contains surface-adherent bacteria.

We then conducted 16S rRNA gene sequencing to define strain-level microbial differences among serially washed and bleached animals. From these data emerged candidate gut-(*Stenotrophomonas* JUb19, *Ochrobactrum* MYb71) and skin-dominant bacteria (*Enterobacter* JUb101, *Sphingobacterium* BIGb0170). Although our two independent sequencing experiments revealed different proportions of microbes (Supplementary Figure 2A,C), both experiments suggest that there are distinct communities in animals with intact microbiomes compared to those without surface microbes (Figure 2C-J). Importantly, the microbiomes from animals with cuticle-adherent microbes are distinct from those of their bacterial lawns (Figure G, I), indicating some level of selection on the surface of the cuticle. We further confirmed the persistence of skin-resident microbes compared to more gut-resident microbes by measuring bacterial growth from skin swabs of animals reared on these species (Figure 3D-G). We hypothesize that multiple species within mCeMbio may also associate with the *C. elegans* cuticle to varying degrees.

None of the mCeMbio bacteria are known to be pathogenic, although the individual isolates demonstrate varying effects on host developmental rates (Dirksen et al. 2020). Given the symbiotic nature of these bacteria and our parallel lines of validation of surface-adherent bacteria, we hypothesized that the mCeMbio community serves a role in cuticle protection against environmental stressors. We conducted a well-established cuticle fragility assay and observed a consistent role of mCeMbio and CBX151, a known cuticle-adherent bacterium, to promote cuticle integrity of the host, which varies across the individual bacterial species in mCeMbio (Figure 4A,B). From these data, we demonstrate the importance of studying community effects, and the effects of single bacterial species on cuticle integrity. Put together, we uncover a protective effect for animals reared on a consortium of ecologically relevant bacteria.

Our data do not differentiate between the combinatorial effects of the mCeMbio bacteria on the skin and in the gut, and the mechanism of this protection requires further examination. Several species of bacteria found in association with *C. elegans* are known to, directly and indirectly, protect against pathogens in the worm intestine (Kohar A. B. Kissoyan et al. 2019). This may provide an indirect mechanism by which gut-dominant bacteria are able to protect the cuticle. Moreover, both known skin-resident bacteria and members of the mCeMbio consortium have been shown to prime the cuticle against other infections (Jonathan Hodgkin et al. 2013). This protection may even be attributed to the potential hydrogen peroxide-degrading capabilities of the mCeMbio consortium, which have been observed in strains of *E. coli* (Schiffer et al. 2021). Importantly, we sought to examine skin-enriched microbes and the mCeMbio consortium as a whole, which involves both bacteria-bacteria and host-bacteria interactions. This is highlighted by greater cuticle resistance animals raised on the mCeMbio community (time to burst = 277 seconds) compared to all but one of the individual mCeMbio members, the gut enriched strain *Ochrobactrum* MYb71 (Figure 4B). This suggests that most single host-bacteria interactions may not fully recapitulate the extent of interactions in our system and demonstrates the importance of studying communities of ecologically relevant microbes.

In an effort to develop *C. elegans* as a model for understanding skin and associated diseases, we must make use of sensitized and protective strains. To do this, we used *C. elegans* mutants with varying levels of disordered cuticle structure and function. We found that mCeMbio and our strains of interest, JUb19 and JUb101, as well as the known skin-adherent microbe CBX151, increase protection in both sensitized and protective mutants (Figure 4C-E, Supplementary Figure 4A-D).

Given the symbiotic nature of the mCeMbio bacteria, which includes 11/12 published CeMbio strains, unique cuticle structures of the nematode skin such as the annuli, alae, or invagination sites, and community effects on promoting barrier function, our data support the existence of and role for a previously unappreciated *C. elegans* skin microbiome. Our work supports the continued efforts to make the study of *C. elegans* more ecologically relevant and suggests that we continue to understand and increase environmental enrichment to improve the relevance of this model system.

## Materials and Methods

### C. elegans strains and maintenance

Strains were ordered from the Caenorhabditis Genetics Center and include the following: wild type N2; CB130 *dpy-8(e130)*; VC2985 *dpy-10(gk3075)* II; CB6627 *srf-3(e2689)* IV; CB6177 *bus-8(e2883)* X. All animals were grown at 20°C under standard conditions to adulthood (Stiernagle 2006). NGM plates were seeded with either *E. coli* OP50 or mCeMbio strains unless noted of a specific bacterium. For all experiments, L4 animals are placed on lawns and removed ∼12 hours after, leaving only offspring behind. Experiments are then performed on the offspring unless otherwise noted.

### Culture and maintenance of mCeMbio bacteria

Individual isolates of “CeMBio” bacteria, referred to as mCeMbio (gift from H. Schulenberg, available now through CGC as CeMbio, see note below in Identification of mCeMbio) and CBX151 (a gift from Marie-Anne Félix) were streaked onto LB agar plates and individual colonies were picked and grown in LB growth media. Individual bacteria (12 mCeMbio strains, CBX151, and OP50) were allowed 2-4 days to grow, shaking at 4 x g at 20°C. Bacteria were maintained prepared to OD_600_ = 0.2 by dilution in additional LB or centrifuging at 1800 x g for 5 minutes to remove excess LB to increase the concentration of slow growing isolates. To make mCeMbio, each of the 12 bacteria were combined at equal ratios. 200 µL of the bacteria were pipetted on 6 cm NGM plates and evenly spread across the agar plate using a sterile L spreader (VWR 490007-358). Bacterial plates were left to dry in a laminar flow hood for 2 days. If not used immediately, seeded plates were stored at 4°C. We maintain glycerol stocks of each bacterium at −80°C and these were used for each new batch of each mCeMbio strain, OP50, or CBX151 bacteria.

### Identification of mCeMbio strain JUb101

The *Lelliottia amnigena,* JUb66, genome sequenced in Houston by Buck Samuel’s group and the JUb66 genome sequenced in Kiel by Heinrich Schulenburg’s group are different. The bacterial sample deposited at the CGC as part of the CeMbio consortium is the JUb66 from the Samuel Lab (JUb66 Houston) and the one from the Schulenburg lab is now known as JUb101. We acquired our strains from the Schulenburg lab. These two genomes have identical 16S rRNA representative sequences and the same number of 16S repeats. A full genome comparison of JUb66 Houston and JUb101 (formerly known as JUb66 Kiel) are only 84% similar. To determine the identity of our CeMbio strain, formerly known as JUb66 Kiel, we used the following PCR primers and standard NEB Taq protocols.

JUb66-2270F TTGCTTCGGGTGGTGGTATC JUb66-2827R CCATGGAACTCACCCCTGTC JUb101-1377F GAGGATGGGTATGCAGACCG JUb101-2375R GGGATCGCAATCTGACGGAT Therefore, we call our mixture mCeMbio, a modified CeMbio consortium.

### C. elegans and bacterial prep for DNA extraction

Animals were reared on mCeMbio bacterial lawns for 48-82 h after hatching. Bacterial plates (n=3 for each group) were flooded with M9 and 30 animals were picked off with a glass pipette into an Eppendorf tube. For unbleached animals, tubes were washed with M9+T (M9+25mM Tetramisole Hydrochloride (Sigma-Aldrich L9756)) 5 times, allowing 4 minutes for animals to sink to the bottom of the tubes between each wash. For bleached animals, tubes were washed with M9+T 2 times and then washed with surface-sterilizing bleaching solution (M9+T+2.5% hypochlorite+1M KOH) for 4 minutes. These tubes were rinsed with M9+T twice for 4 minutes each. After washes, animals were allowed to sink to the bottom of the tube. All but 100 μL of media was removed in the final wash and the tubes were frozen on dry ice and stored at −80°C. For “lawn” plates, 1 mL of M9 was added to the plate and the bacterial lawns were scraped with a sterile L spreader (VWR 490007-358). Bacterial slurry was pipetted and centrifuged at 13000 x g for 4 minutes. All but 100 μL was removed and the pellet was frozen on dry ice and stored at −80°C. DNA was extracted following the standard NucleoSpin Tissue Kit (Macherey-Nagel 740952) protocol for animal tissue with 1 hour of incubation for the first lysis step. DNA was stored at −20°C.

### 16S rRNA amplicon sequencing and analysis

For each experiment, an amplicon library targeting the V4-V5 hypervariable region of the 16S gene was generated via one-step PCR using the Azura TruFi DNA Polymerase PCR kit and the 515-F (GTGYCAGCMGCCGCGGTAA) and 926R (CCGYCAATTYMTTTRAGTTT) primer set was used (Parada, Needham, and Fuhrman 2016). Each reaction was performed with an initial denaturing step at 95°C for 1 minute followed by 30 cycles of 95°C for 15 seconds, 56°C for 15 seconds, and 72°C for 30 seconds. 2.5 µL of each PCR reaction was run on a 1.8% agarose gel to confirm amplification. PCR products were purified using Beckman Coulter AMPure XP following the standard 1x PCR clean-up protocol. PCR quantification was performed in duplicate using Invitrogen Quant-iT PicoGreen dsDNA Assay kit. Samples were then pooled in equal proportions (20 ng DNA per sample) followed by another AMPure XP PCR purification. The final libraries were evaluated on an Agilent 2200 TapeStation (D1000 ScreenTape), quantified with Qubit HS dsDNA kit, and each sequenced at the University of California, San Diego IGM Genomics Center on a single Illumina MiSeq lane with a 25% PhiX spike-in.

Amplicons were analyzed with QIIME2 v2019.10 (Bolyen et al. 2019). Briefly, demultiplexed paired-end reads were trimmed to remove adapter and primer sequences with cutadapt (Martin 2011). Trimmed reads were then denoised with DADA2 to produce amplicon sequence variants (ASVs) (Callahan et al. 2016). Taxonomic annotation of ASVs was conducted with the q2-feature-classifier classify-sklearn naïve-bayes classifier (Pedregosa et al. 2011; Bokulich et al. 2018) both the genome-derived reference 16S sequences of CeMbio strains (Dirksen et al. 2020) and the SILVA database (v138) (Pruesse et al. 2007). ASVs were then aggregated by bacterial strain and reads were normalized by the 16S gene copy numbers estimated by their genome assemblies(Dirksen et al. 2020). Bray-Curtis dissimilarities, Shannon Indices, and tests of statistical significance were then calculated with the QIIME2 diversity plugin on the normalized relative abundance count data (Kruskal and Wallis 1952; Anderson 2001). Raw sequence data are deposited in the Sequence Read Archive (SRA) under the BioProject accession <u>PRJNA979901</u>. Data are sorted into experiment 1 and experiment 2 “combine”, in which all wildtype, hermaphrodite samples, regardless of date of extraction or age, were included if they met the category “bleached” or “unbleached” (serially washed). These data are displayed in Figure 2 and sorted by experiment. Supplementary Figure 2 displays data from combining experiment 1 and experiment 2, but only the samples in which the bleached and unbleached (serially washed) samples were prepared as a matched set on the same day and at day 1 of adulthood.

### Skin-swabbing with 96-well plate assay

To make the micro swab, a glass capillary tube (World Precision Instrument Thin Wall Glass Capillaries-TW100-4 or similar) was pulled apart over a small flame into two pieces. Heat was used to create a small closed bulb at the pulled end of each piece, ensuring there were no crevices in which bacteria or liquids may accumulate, and allowed the glass to cool. Then using 600-grain sandpaper, the bulb was lightly scored to make a textured surface (∼10 small circles on each side and tip). To sterilize, the swab is dipped in 70% ethanol and air dried.

To swab a lawn, the micro-swab is dipped into the lawn on which the animals were reared and then immediately swirled in the appropriate LB-containing wells. To swab an animal with the condition “crawl”, a single animal is placed on an empty NGM agar plate and allowed to crawl for 30 seconds. Using a dissecting microscope, the animal was gently swabbed using the bulbous end from head to tail 10 times. The bulb was then dipped and swirled around in 10 circles into 200 μL of LB in a 96 well plate (CoStar 96 Flat Black). Animals that were washed once “1x wash” were washed in M9+T and allowed to sink to the bottom of a tube. This was repeated two more times for “3x wash”. For animals in the “bleach” group, they were first washed with M9+T 3 times, then once with surface-sterilizing bleaching solution (M9+T+2.5% hypochlorite+1M KOH), and once more with M9+T. Once washed or bleached, the animals were then pipetted onto a sterile NGM plate in a small volume and allowed to crawl out of the drop before being swabbed. Once all animals were swabbed, the OD_600_ was measured using a Molecular Devices SpectraMaxPlus with absorbance = 600 nm, manually taking measurements during standard work hours. To generate the curves in Figures 1D, F and Figures 2B, D, F, the individual replicate OD_600_ values at each time point were averaged in each condition. Data above the initial maxima were removed. A simple linear regression analysis was applied to fit the data with unknowns interpolated from a standard curve.

### Colony Forming Unit Assay

Animals were reared to day 1 of adulthood on chosen bacteria. 50 adult worms were transferred to a 2 mL tube containing 100 μL M9 and 10-20 1 mm zirconium beads (Beadbug Z763780). Each tube of worms was washed 3 times for 3 minutes each with 750 μL M9+Tetramisole (25mM), removing all but 100 μL liquid with a new pipette tip each time, allowing animals to sink by gravity. We separated half of the tubes to be “bleached” groups, which are subjected to a 4-minute wash with surface-sterilizing bleaching solution (M9+T+2.5% hypochlorite+1M KOH). Bleached animals are washed for 3 minutes x 2 using M9-T to remove the bleach. On the final wash, 100 μL of the supernatant from each bleached tube is removed. Unbleached samples are washed for 3 minutes x 2 using M9-T and 100 μL of the supernatant is removed for each unbleached tube. Each sample is homogenized using a Fisher Mixer Mill at Level 4 for 30 seconds for 4 cycles, totaling 2 minutes of bead beating. The homogenate was then serially diluted (1 in 100, 1000, 10000, 100000 for each tube of 50 worms) in M9. 100 μL of each dilution was plated and spread with a sterile L spreader on LB agar plates. The previously removed supernatants were also diluted in M9 and plated. Plates were dried and left to grow for 48 hours. Finally, bacterial colonies were counted to quantify the original bacterial colony counts in each sample (unbleached, bleached, and supernatant of both bleached and unbleached samples).

### Cuticle Fragility Assay

15 μL of strong bleach solution (1M KOH and ∼5% hypochlorite) is pipetted onto a single day 1 adult animal after it has crawled on a clean NGM plate for ∼10 seconds. Under a dissecting microscope, the time to burst open (cuticle loses integrity) is recorded.

### Standard Statistical Analyses

Figures 1, 3-5 and Supplementary Figures 1, 3-5 and their statistical analyses were completed in Graph Pad Prism. Ungrouped 2-sample t-tests were used when determining the statistical significance during comparisons of 2 groups. With 3 groups and more, an ANOVA test for significance values was performed. When appropriate, corrections were performed for multiple comparisons or unequal variances. Any *P* value smaller than 0.05 was considered significant. The statistical tests used in Figure 2 are described under “16S rRNA Amplicon Sequencing and Analysis”.

## Data availability

Raw sequencing data for the 16S sequences are deposited at the National Center for Biotechnology Information Sequence Read Archive under the BioProject Accession <u>PRJNA979901</u>. All other data, including raw microscopy images, are available upon request.

## Acknowledgments.

We would like to thank the Chalasani lab members, especially Kirthi Reddy, for their support and feedback on this nascent project. We are grateful to the Schulenberg, Samuel, & Dierkling lab members for access to mCeMbio strains and discussion. We thank Dr. Andrew Allen for the use of his lab space at the J. Craig Venter Institute to prepare DNA for 16S sequencing. The idea to develop a micro-swab was initiated by Dr. Brooke Pickett, NBH’s microbiology professor at UCSD. All *C. elegans* strains used were provided by the CGC, which is funded by NIH Office of Research Infrastructure Programs (P40 OD010440). The Illumina MiSeq 16S Sequencing was conducted at the IGM Genomics Center, University of California, San Diego, La Jolla, CA. MAM was supported by NSF PRFB #2011023, with supplemental funding provided by the Hoffman Foundation. NBH was supported by the UCSD Faculty Mentoring Program, the Summer URS Ledell Family Research Scholarship for Science and Engineering, with technical and professional support provided by UCSD’s Research Scholarship Coordinator, Dr. Sophia Tsai Neri. BSS was supported with NIH grant DP2DK116645, NASA grant 80NSSC22K0250, and JGI/DOE grant CSP-503338. SHC was supported by an NIH grant R01MH096881. Figures 1A,C, Supplementary Figure 2D, and Figure 4A were created by the authors with BioRender.com.

## Supplementary Figure Legends

**Figure S1: Growth curves of swabbed bacteria reveal mCeMbio, but not OP50 grows after washing.** Raw OD_600_ of swabbed bacteria from animals reared on OP50 (A,C) and mCeMbio (B,D) during 48 hours of growth. Each curve is the growth from swabs of a single worm. Data, along with experiments in Figure 1 D and F are used to generate Area Under Curve for Figure 1E and 1G.

**Figure S2: Combined, matched data from both experiments reveals consistent differences between washed and bleached animals.** (A) Principal coordinate analysis of Bray-Curtis dissimilarities for matched bleach/washed sets in experiments 1 and 2. B) Bray-Curtis dissimilarities as distance from the bleached group of matched bleach/washed sets in combined experiments 1 and 2 with pairwise PERMANOVA q-values. C) Bray-Curtis dissimilarities as distance from all groups in experiment 1 compared to all groups in experiment 2 with pairwise PERMANOVA q-values.

**Figure S3: CBX151 and JUb101 remain on C. elegans cuticle after washes, while JUb19 does not.** Raw OD_600_ of swabbed bacteria from animals reared on CBX151 (A, B), JUb101 (C, D), and JUb19 (E, F) during 72 hours of growth with n≥5 animals for each condition. Each curve is the growth from a swab of a single worm. These data, together with Figure 3B, D, and F are used to generate Area Under Curves for Figure 3C, E, and G.

**Figure S4: CBX151, mCeMbio and some individual species protect wildtype and mutant C. elegans from harsh bleach.** A) Average time to burst for N2, *bus-8, dpy-8, srf-3,* and *dpy-10* animals raised on OP50 or mCeMbio at days 1, 2, and 3 post adulthood. N=3, n=9 animals in experiment. Mean with standard deviation. B-D) Time to burst normalized to OP50 control for N2 (B), *bus-8* (C), and *dpy-10* (D) animals raised on OP50, CBX151, mCeMbio, JUb19, or JUb101, mean with standard deviation. Each dot is an individual animal, normalized to the average of the OP50 controls on the day it was tested. N=3 experiments with n = 9 animals in each experiment A) Two-Way ANOVA with significance between the age/strain, bacteria, and the interaction between them with Šídák’s multiple comparisons test between OP50 and mCeMbio for each age/genotype. B-D) Brown-Forsythe One-Way ANOVA with Dunnett’s T3 Multiple Comparisons Test * p<0.05, ** p<0.01, *** p<0.001, **** p<0.0001.

**Supplementary Table 1:** *C. elegans* strains used in this study.

| ID | gene (allele) | Phenotype |
| --- | --- | --- |
| N2 |  | Wildtype |
| CB130 | *dpy-8* (*e130*) | no annuli |
| VC2985 | *dpy-10*(*gk3075*) | no annuli |
| CB6627 | *srf-3*(*e2689*) | resistant to biofilms |
| CB6177 | *bus-8*(*e2883*) | disordered epidermal morphogenesis |

## Notes

https://www.ncbi.nlm.nih.gov/sra/PRJNA979901

### Competing Interest Statement

The authors have declared no competing interest.

