## SupplementaryFigures1-4 for "Identification and characterization of a skin microbiome on *Caenorhabditis elegans* suggests environmental microbes confer cuticle protection"

Supplementary Figure 1

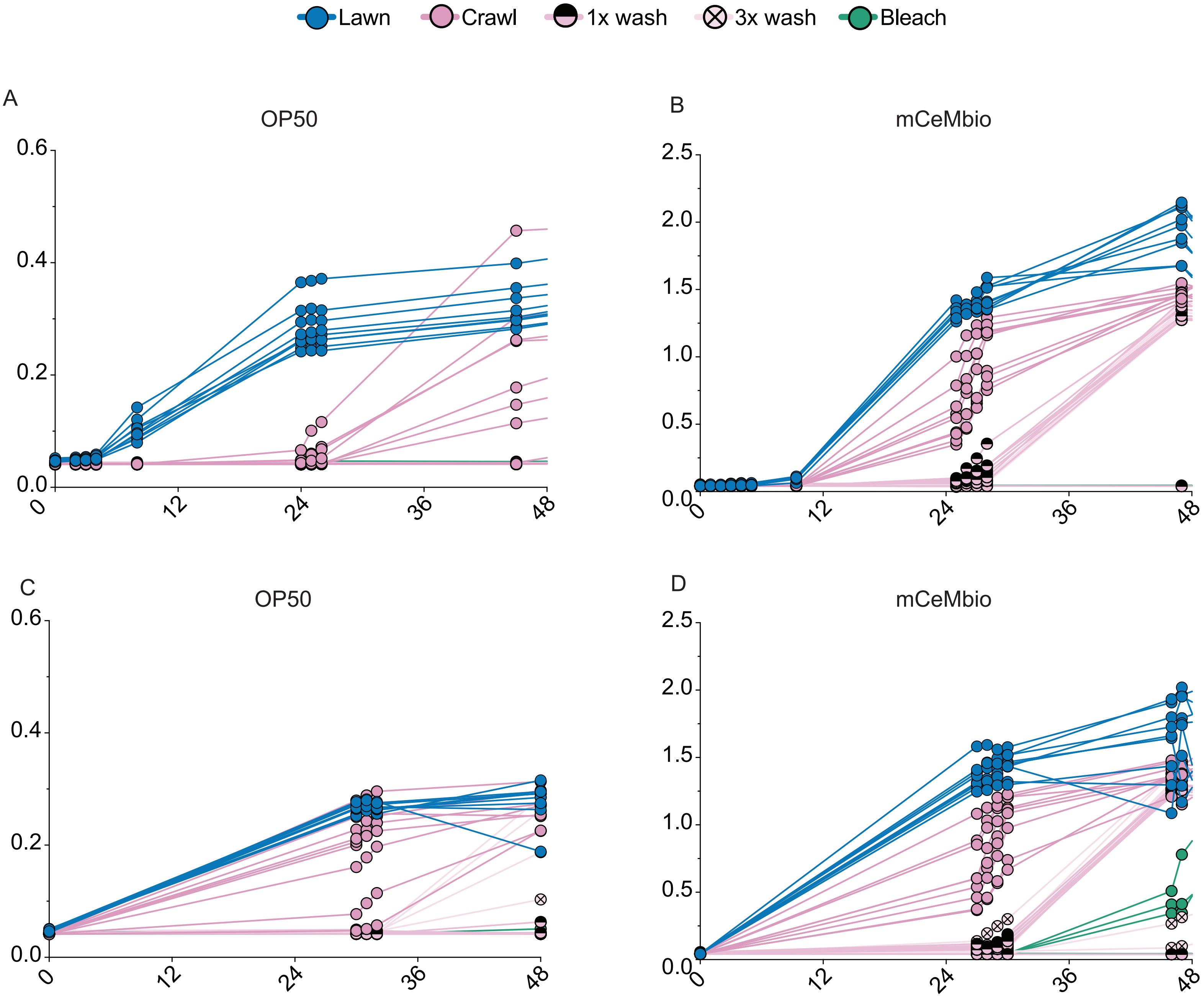

Supplementary Figure 2

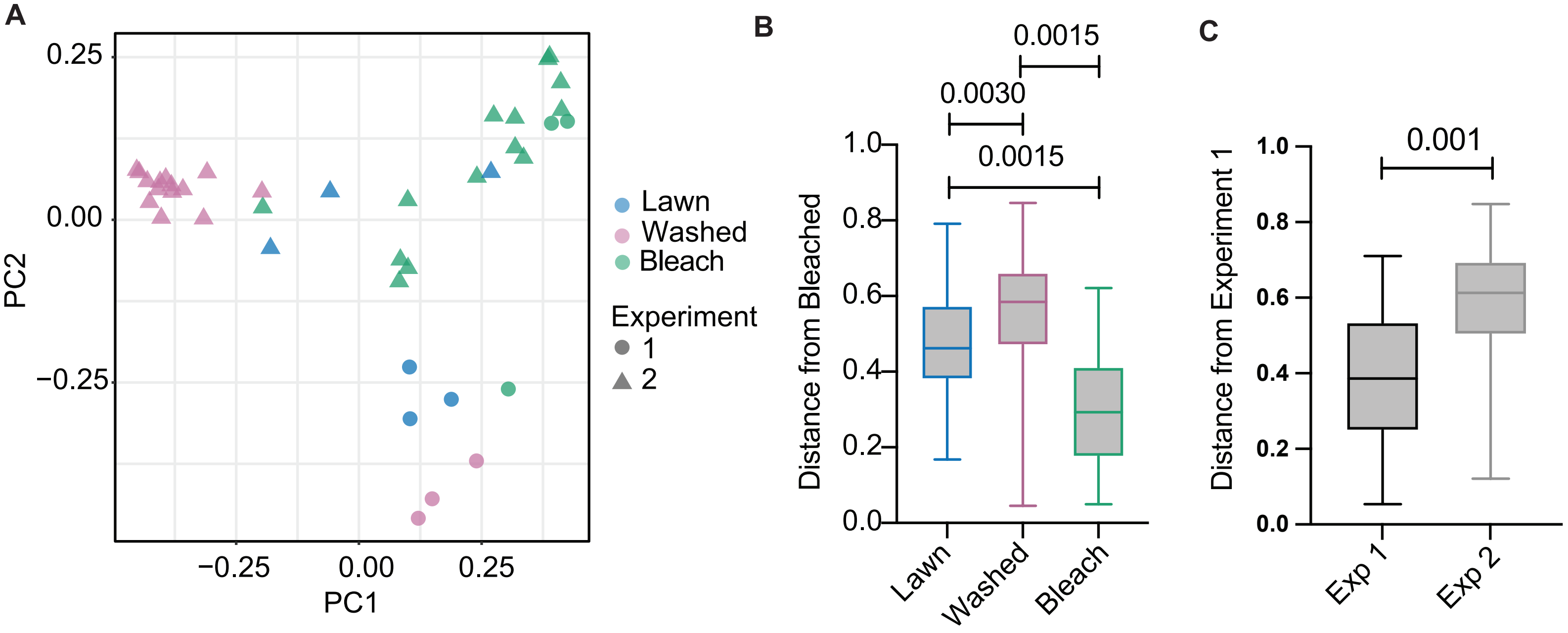

Supplementary Figure 3

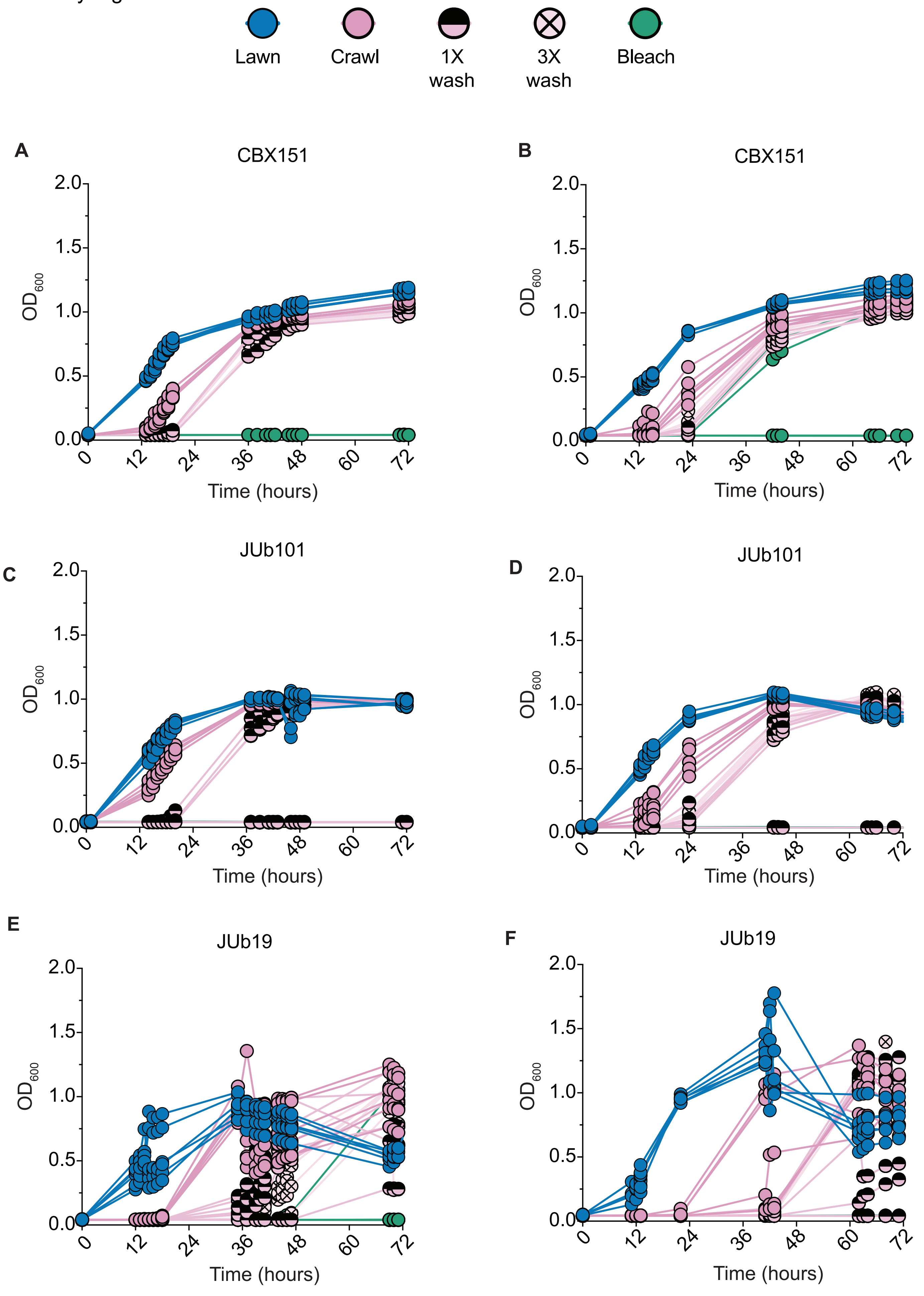

Supplementary Figure 4

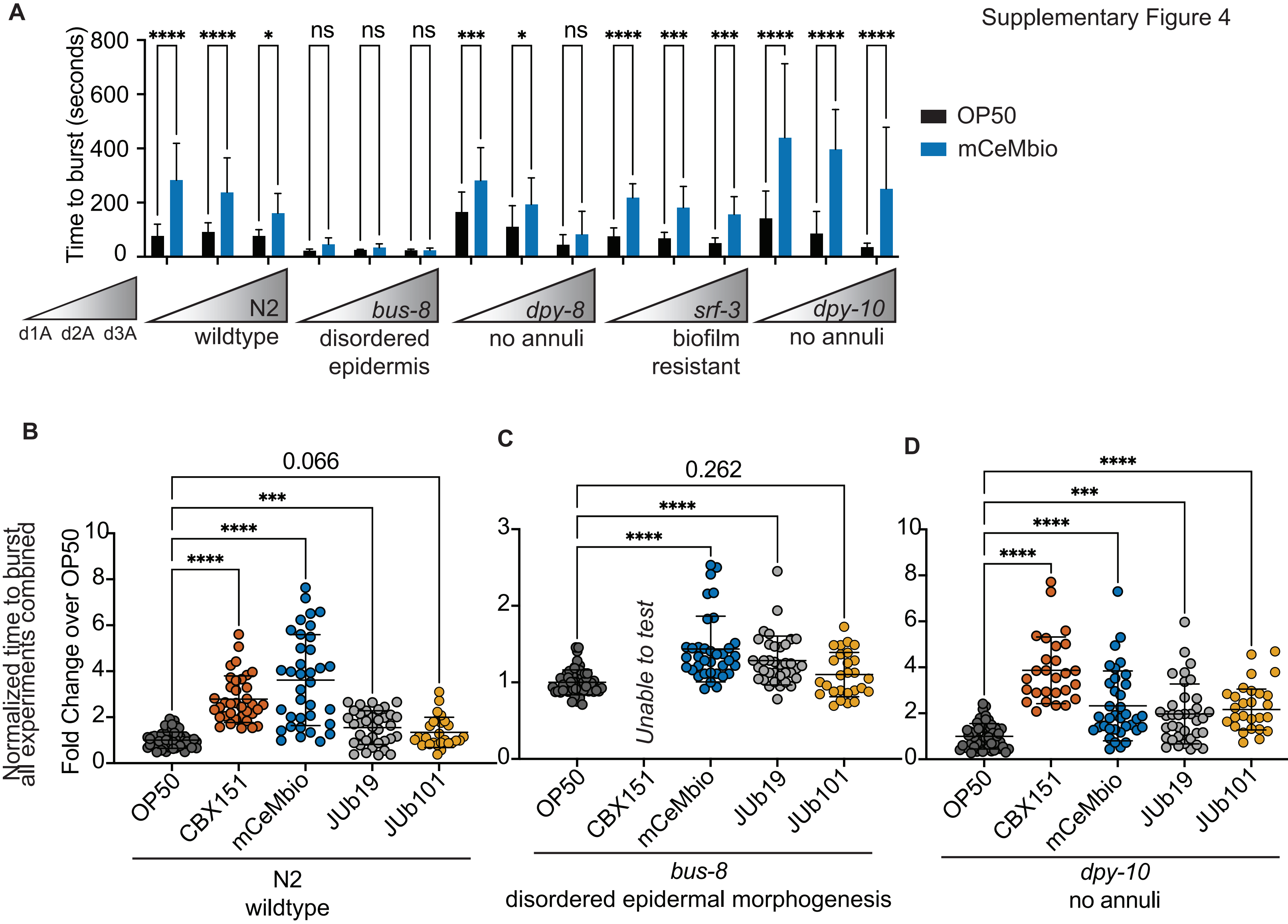

Supplementary Figure Legends:

**Figure S1: Growth curves of swabbed bacteria reveal mCeMbio, but not OP50 grows after washing.** Raw $OD_{600}$ of swabbed bacteria from animals reared on OP50 (A,C) and mCeMbio (B,D) during 48 hours of growth. Each curve is the growth from swabs of a single worm. Data, along with experiments in Figure 1 D and F are used to generate Area Under Curve for Figure 1E and 1G.

**Figure S2: Combined, matched data from both experiments reveals consistent differences between washed and bleached animals.** (A) Principal coordinate analysis of Bray-Curtis dissimilarities for matched bleach/washed sets in experiments 1 and 2. B) Bray-Curtis dissimilarities as distance from the bleached group of matched bleach/washed sets in combined experiments 1 and 2 with pairwise PERMANOVA q-values. C) Bray-Curtis dissimilarities as distance from all groups in experiment 1 compared to all groups in experiment 2 with pairwise PERMANOVA q-values.

**Figure S3: CBX151 and JUb101 remain on C. elegans cuticle after washes, while JUb19 does not.** Raw $OD_{600}$ of swabbed bacteria from animals reared on CBX151 (A, B), JUb101 (C, D), and JUb19 (E, F) during 72 hours of growth with $n \geq 5$ animals for each condition. Each curve is the growth from a swab of a single worm. These data, together with Figure 3B, D, and F are used to generate Area Under Curves for Figure 3C, E, and G.

**Figure S4: CBX151, mCeMbio and some individual species protect wildtype and mutant C. elegans from harsh bleach.** A) Average time to burst for N2, *bus-8, dpy-8, srf-3,* and *dpy-10* animals raised on OP50 or mCeMbio at days 1, 2, and 3 post adulthood. N=3, n=9 animals in experiment. Mean with standard deviation. B-D) Time to burst normalized to OP50 control for N2 (B), *bus-8* (C), and *dpy-10* (D) animals raised on OP50, CBX151, mCeMbio, JUb19, or JUb101, mean with standard deviation. Each dot is an individual animal, normalized to the average of the OP50 controls on the day it was tested. N=3 experiments with n = 9 animals in each experiment A) Two-Way ANOVA with significance between the age/strain, bacteria, and the interaction between them with Šídák's multiple comparisons test between OP50 and mCeMbio for each age/genotype. B-D) Brown-Forsythe One-Way ANOVA with Dunnett's T3 Multiple Comparisons Test * $p<0.05$, ** $p<0.01$, *** $p<0.001$, **** $p<0.0001$.
